# Tracking a major evolutionary transition to superorganismality

**DOI:** 10.64898/2026.08.23.746534

**Authors:** Bitao Qiu, Shuangrong Li, Zewen Zhou, Joh Henschel, Robert Hanus, Bao Jia, Qionghua Gao, Judith Korb

## Abstract

Major transitions in evolution are associated with the loss of independent reproduction by formerly autonomous units. Termites provide a powerful system for studying this process because they exhibit diverse social systems in which workers’ developmental and reproductive potential declines with increasing colony-level organismality. However, the evolutionary sequence and developmental genetic basis of these transitions remain unresolved. Here, using comparative developmental transcriptomics across seven termite species that differ in workers’ reproductive potential, we reconstructed the evolutionary history of termite social systems. We found that linear caste development, in which workers retain full reproductive potential, represents the ancestral state of termites. Bifurcated caste development, in which workers partially lose reproductive potential early in development, evolved independently multiple times, with two origins subsequently giving rise to superorganisms with unipotent, sterile workers. Ancestral gene regulatory network (GRN) reconstruction revealed that linear caste development evolved through retention of a juvenile-like worker state and co-option of a conserved developmental GRN characterizing hemimetabolous insect nymphal development, in which juvenile hormone, ecdysone and TGF-β signaling pathways play central roles. The convergent evolution of bifurcated caste development repeatedly co-opted the GRN underlying linear caste development, heterochronically shifting its activity to earlier developmental stages. Finally, we found that the evolution of termite superorganisms involved somatization of the worker caste and co-option of a conserved endocrine GRN for terminal differentiation. Together, these findings uncovered repeated routes to reduced workers’ reproductive potential through GRN co-option and highlight striking parallels between superorganism evolution in social insects and organismal evolution in metazoans.

## Introduction

The evolution of life is characterized by major evolutionary transitions in individuality (METs) that led to increasing complexity, from prokaryotes to eukaryotes, from unicellular to multicellular organisms (Szathmary and Smith 1995; West et al. 2015). Insect sociality, as classically found in termites and social Hymenoptera (ants and some bees and wasps), has been regarded as the latest METs, adding a new level above the organism: the superorganism(West et al. 2015; P. Kennedy et al. 2017; Boomsma and Gawne 2018; Lohmar et al. 2026). During METs, pre-existing lower-level evolutionary units overcome the omnipresent Darwinian forces of competition and conflict, evolving cooperation to the point that they form a new higher-level evolutionary unit, which becomes the new main target of selection (Szathmary and Smith 1995). The evolution of multicellularity, as in metazoa, and insect superorganismality share common traits, including high relatedness and reproductive division of labor among lower-level units forming a higher level (West et al. 2015; Howe et al. 2022). In metazoa and social insects, respectively, a few cells/individuals specialise on reproduction (the germline/ queens and sometimes kings; hereafter we refer to queens only), while the large majority perform other tasks, refraining from reproduction (somatic cells/workers and sometimes soldiers; hereafter we refer to workers only) (West et al. 2015). This reproductive altruisms of somatic cells and workers presented an evolutionary puzzle which can ultimately be explained by kin selection theory (Hamilton 1964a, 1964b). Mechanistically, it is achieved in metazoans through germline/soma segregation during development, which prevents somatic cells from becoming germline and creates Weismann’s barrier, eliminating somatic cells from obtaining direct fitness (Buss 1987; C. G. M. Extavour 2007).

In many social insect species, workers are not sterile but can reproduce to varying degrees. Three categories can be distinguished (i) totipotent worker, which retain the full capacity to reproduce, including becoming dispersing reproductives, (ii) pluripotent workers, which can still reproduce but cannot become dispersing reproductives, and (iii) unipotent workers, which are functionally sterile, similar to somatic cells in metazoans (Lohmar et al. 2026). Along these categories, declining workers’ reproductive potential result in decreasing within-colony conflict and increasingly aligned evolutionary fitness interests, giving rise to a superorganismality gradient. At its extreme end, in species with unipotent sterile workers the colony becomes the main target of selection and adaptation (Gardner and Grafen 2009). Notably, also in metazoans, similar diversity exists. As first pointed out by Buss and increasingly recognized since (Buss 1983; C. G. M. Extavour 2007; Devlin, Ganley, and Takeuchi 2023; Lohmar et al. 2026), many animals, especially within the nonbilaterian lineages, retain lifetime toti- or pluripotent cells that can subsequently give rise to germ cells. This variation in cell totipotency is generally associated with the developmental timing of germline/soma segregation and possibly the degree of organismal complexity (Buss 1987; Devlin, Ganley, and Takeuchi 2023; Howe et al. 2022). These striking analogies between metazoans and insect superorganismality ask for a better understanding of the evolutionary history and the mechanisms underlying the METs to superorganismality will shed light on shared principles. While the evolution history and the developmental mechanisms for germline-soma segregation in metazoan are well studied, comparable work in social insects remains heavily biased toward Hymenoptera (Evans and Wheeler 1999; Berens, Hunt, and Toth 2014; Warner et al. 2019; Qiu et al. 2022), and none has yet tracked the full transitions to superorganismality within a single lineage.

Among social insects, termites provide a particularly good example of the superorganismality gradient as toti-, pluri- and unipotent systems can clearly be distinguished (J. Korb 2025). Species with totipotent workers — sometimes called ‘pseudergates’ (false workers) — typically have a have a linear development from larval (wingless) via nymphal (with wing buds) instars into winged dispersing reproductives (**Figure 1**). The wingless stages of termites are unusual because as hemimetabolous insects their immatures should be nymphs. In contrast, workers of other species — sometimes called ‘true workers’ — are either pluripotent (i.e. they cannot develop into dispersing reproductives but can still produce within the nest as neotenic reproductives) or unipotent, when they are functionally sterile (Figure 1). Analogous to germline/soma segregation in metazoans, this differentiation of true workers is achieved through a bifurcated caste development with an apterous (wingless) line, from which (mainly) workers derive, and a nymphal line from which dispersing reproductives develop (for more details see Figure 1). For simplicity, we will call apterous working instars ‘workers’ and nymphal instars ‘nymph’ in the following, although both can perform worker tasks (Roisin and Korb 2011; Judith Korb 2025). Because linear and bifurcated caste development programs are found in both basal and derived termite lineages, the ancestral state of termite workers has been debated for more than 40 years and both developmental programs are thought to have evolved independently multiple times (Watson and Sewell 1985; Noirot and Pasteels 1987; Thompson et al. 2004; Grandcolas and D’Haese 2004; Legendre et al. 2008; Hellemans et al. 2024). This variation in workers’ reproductive potential therefore provides a unique opportunity to track the full transition to superorganismality and the underlying evolutionary and developmental principles.

**Figure 1.**
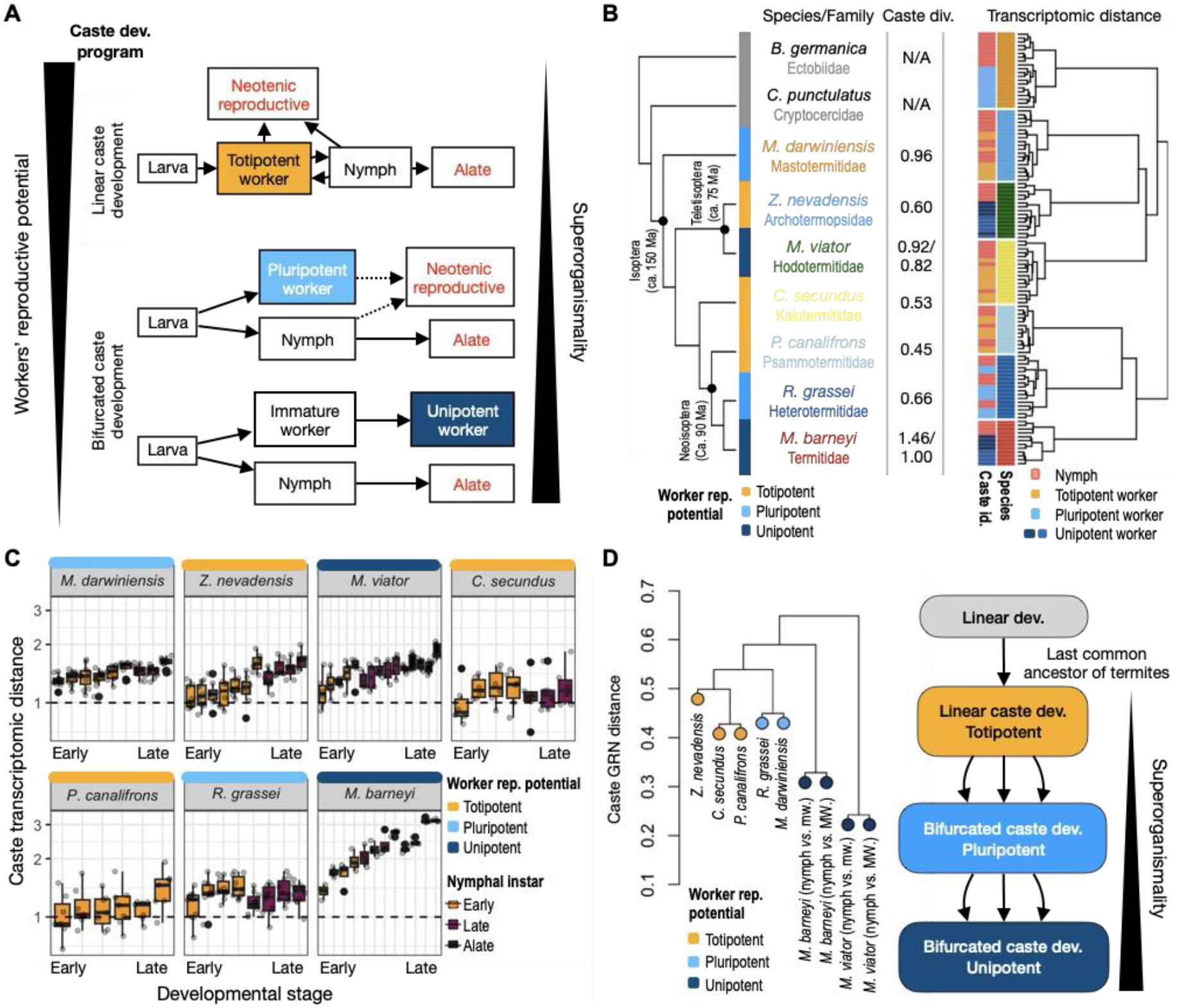
Comparative developmental transcriptomes reconstruct the evolutionary history of the termite caste developmental program. (A) Termite caste developmental programs can be classified as linear or bifurcated caste development and their workers’ reproductive potential declines with increasing colony-level (super-)organismality. For simplicity, soldiers are omitted. (B) Phylogenetic relationships, workers’ reproductive potential, and between-caste transcriptomic divergence of the studied species (left), together with hierarchical clustering based on late-instar transcriptomes (right). In the dendrogram, samples from the same species are indicated by colours matching those in the left panel. (C) Boxplots of normalized between-caste transcriptomic distance across developmental stages and termite species. Normalized between-caste distance was calculated as the log ratio of the between- caste transcriptomic distance to the within-worker-caste transcriptomic distance. (D) Hierarchical clustering based on distances among caste-biased expression profiles between nymphs and workers, representing the caste developmental GRNs, across seven termite species (left). The analysis included one-to-one termite orthologs showing significant caste-biased expression in at least one species (*n_genes_*= 3,527). For *M. viator* and *M. barneyi*, both nymph–minor worker (mw) and nymph–major worker (MW) contrasts were included. Clustering of caste developmental GRN supports linear caste development with totipotent workers as the ancestral state of termites and bifurcated caste development evolved convergently (right).

We sequenced genomes and generated developmental transcriptomes across the termite phylogeny, including several species with toti-, pluri-, and unipotent workers to reconstruct the gene regulatory networks (GRNs) underlying their caste developmental program and identify homology and convergence across several independent transitions. Our study clearly showed an evolutionary history of decreasing workers’ reproductive potential across the termite superorganismality gradient. The mechanisms underlying these transitions involve co-option of ancient developmental GRNs, coupled with heterochronic shifts of their expression and the retention of a juvenile developmental state in worker castes. Finally, transitions to unipotent, sterile workers were achieved through co-option of an ancestral adult differentiation GRN, terminally differentiating workers into the functional soma of the colony.

## Results

### Developmental transcriptomes mirror termite phylogeny and workers’ reproductive potential

We generated caste-specific developmental transcriptomes from head–prothorax (n = 215) across seven termite species spanning the full gradient of superorganismality and workers’ reproductive potential across the termite phylogeny. These comprise *Mastotermes darwiniensis* (pluripotent worker), *Zootermopsis nevadensis* (totipotent worker), *Microhodotermes viator* (unipotent worker), *Cryptotermes secundus* (totipotent worker), *Prorhinotermes canalifrons* (totipotent worker), *Reticulitermes grassei* (pluripotent worker), and *Macrotermes barneyi* (unipotent worker) (**Figure 1B**) (**Supplementary Table 1**) (**Method**). We additionally used PacBio long-read sequencing to generate high-quality genome assemblies for *P. canalifrons* and *M. viator*, the latter represent the first genome of a Hodotermitidae (**Method**).

We first examined the overall transcriptomic similarity among termite species. Regardless of caste developmental program, transcriptomic samples clustered first by species, then by family/clade. The dendrogram of termite transcriptomes mirrored the phylogeny of termites, with samples from Neoisoptera (*P. canalifrons, R. grassei*, and *M. barneyi*) clustering together and samples from *M. darwiniensis* separated from Teletisoptera (*Z. nevadensis* and *M. viator*) (**Figure 1B**). Principal component analysis (PCA) showed a similar pattern, with the first PC separating Neoisopteran samples from other termite species and the second PC separating *M. darwiniensis* from the Teletisoptera (**Supplementary Figure 1**), indicating that the evolution of termite transcriptomes is largely driven by lineage-specific organ/tissue gene expression, as has been found in other taxa (Brawand et al. 2011; Qiu et al. 2018).

Within each species, samples clustered primarily by developmental stage and secondarily by caste (**Supplementary Figure 1**). Consistent with the expectation that caste developmental trajectories become progressively canalized during development (Qiu et al. 2022), both the between-caste transcriptomic distance and the number of caste differentially expressed genes (caste DEGs) increased with developmental stage (**Figure 1C**; **Supplementary Figure 2**; Methods), showing that caste differences become more pronounced at later stages. Across species, this developmental pattern scaled with the degree of superorganismality: caste divergence at later developmental stages (measured by the standardized effect size of between-caste differences; Methods) was positively correlated with superorganismality (r = 0.87, p = 0.01), being lowest in species with totipotent workers (*C. secundus*, *Z. nevadensis*, and *P. canalifrons*) and highest in species with sterile unipotent workers (*M. viator* and *M. barneyi*) (**Figure 1B**). Consistently, the number of caste DEGs and the number of worker-specific genes also increased with superorganismality (**Supplementary Figure 2**). Together, these results indicate that, in contrast to overall transcriptomic profiles that largely reflect phylogeny, caste canalization increases as workers’ reproductive potential declines and reaches its strongest form in superorganisms.

### Comparative transcriptomics reconstructs the ancestral state of termite workers

To reconstruct the evolutionary history of the caste developmental program in termites—and in particular to test whether the linear caste development with totipotent workers represents the ancestral state—we identified caste DEGs in each species and compared the caste-biased expression profiles between nymphs and workers across the seven termites (*n* _orthologs_ = 3527) (Method). We reasoned that homologous developmental programs should be underpinned by homologous GRNs for caste differentiation, thereby retaining similar caste-biased expression signatures, whereas convergently evolved developmental programs are expected to exhibit more divergent expression patterns (Wagner 2007; Qiu et al. 2018; Feigin et al. 2023).

Hierarchical clustering and PCA of caste-biased expression profiles showed that termites with linear caste development—*Z. nevadensis*, *C. secundus*, and *P. canalifrons*—clustered together in a pattern that mirrored their phylogeny, as expected if their caste GRNs are homologous. By contrast, termites with bifurcated caste development—*M. darwiniensis*, *M. viator*, *R. grassei*, and *M. barneyi*—showed more divergent caste-biased expression profiles that did not follow phylogeny, supporting convergent evolution (**Figure 1D; Supplementary Figure 3**). Notably, the caste-biased expression profiles of *M. viator* and *M. barneyi* were distinct from those of the other termite species, consistent with unipotent workers representing a more derived state. Collectively, these results support a scenario in which the linear caste development with totipotent workers represents the ancestral state of termites, whereas bifurcated caste development with either pluripotent or unipotent workers evolved at least three times convergently.

### Totipotent workers evolved through juvenile-state retention via developmental GRN co-option

To identify the GRN underlying the ancestral state of termites, we reconstructed the ancestral caste- bias expression profile (hereafter ancestral caste DEGs) across *Z. nevadensis*, *C. secundus*, and *P. canalifrons* using ancestral state reconstruction (**Method**). Consistent with the morphological differences between nymphs and totipotent workers, the top ancestral caste DEGs were enriched for genes involved in wing and eye development (**Supplementary Table 2**). For example, vestigial (*vg*), apterous (*ap*), and bifid (*bi*)— key transcriptional regulators of wing development (Tripathi and Irvine 2021) — as well as Histamine-gated chloride channel subunit 1 (*HisCl1*), transient receptor potential- like (*trpl*), and many other genes involved in eye development and pigmentation, were more highly expressed in nymphs than in totipotent workers (**Figure 2A**).

**Figure 2.**
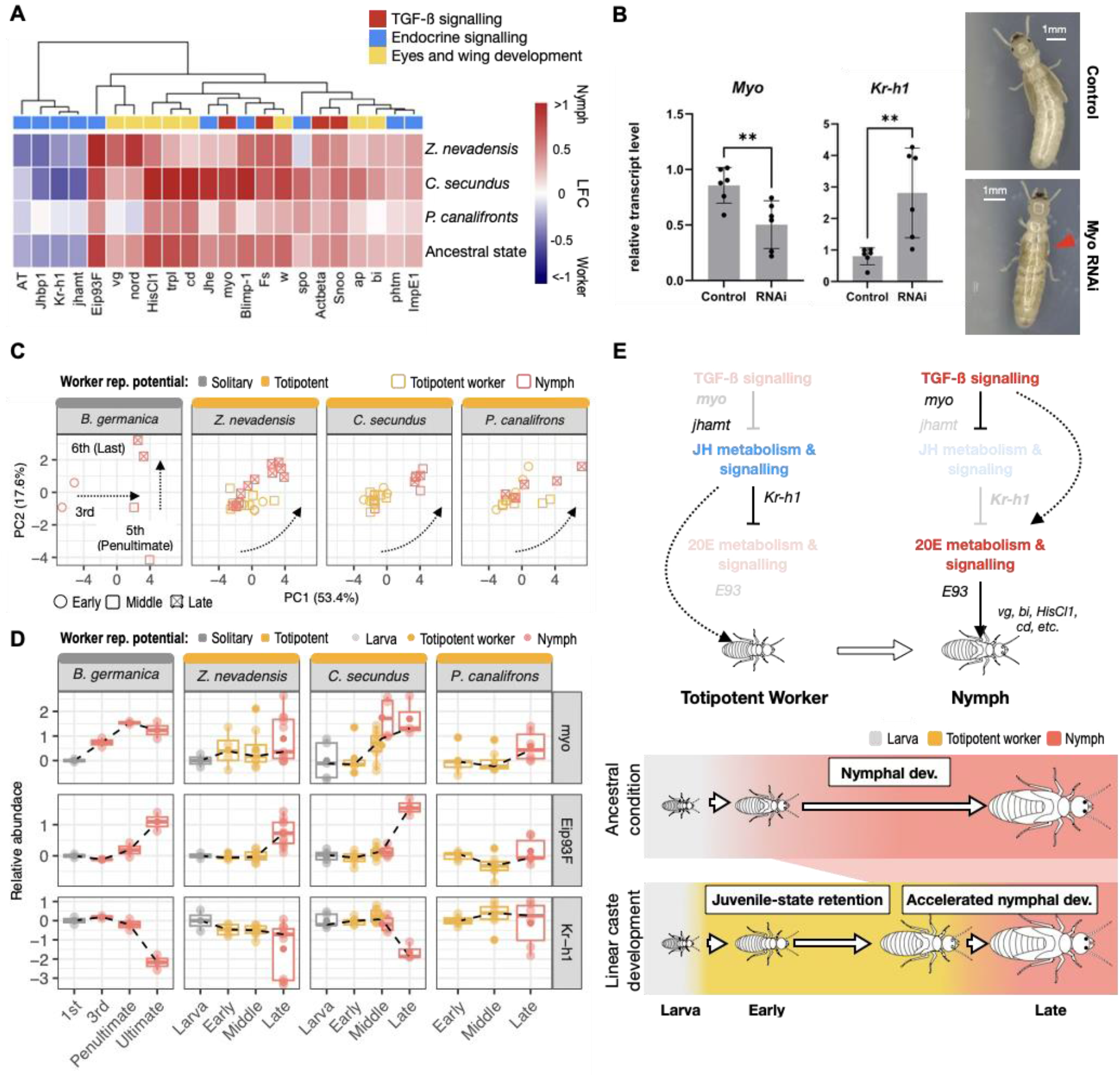
Ancestral state reconstruction reveals the developmental origin of totipotent workers in termites. (A) Heatmap showing caste-biased expression of ancestral caste DEGs associated with key developmental processes across the three termite species with totipotent workers and in the reconstructed ancestral state. The complete gene list is provided in **Supplementary Table 2**. Colours indicate log2 fold changes (LFC) in expression between castes, with red denoting nymph-biased expression and blue denoting worker-biased expression. (B) Relative transcript abundance of *myo* and *Kr-h1* 48 hours after dsMyo injection in *Cryptotermes declivis* nymphs (left), with corresponding post-moult phenotypes of *myo* RNAi knockdown and control individuals (right). Red arrowhead indicates reduced wing buds in *myo* RNAi-treated nymphs. (C) PCA of *Blattella germanica* developmental transcriptomes based on the top 250 ancestral termite caste DEGs, together with the projection of termite transcriptomes into the *B. germanica*-trained PC space. Dashed arrows indicate developmental trajectories. (D) Developmental expression dynamics of the endocrine marker *Kr-h1* and *Eip93F* genes as well as the TGF-β superfamily signalling gene *myo* across termites with totipotent workers and *B. germanica*. Expression levels are shown relative to the earliest developmental stage in each species, as log2 fold change. Dashed lines indicate expression trends based on mean values. (E) Inferred key gene regulatory interactions underlying linear caste development, based on the conserved hemimetabolous developmental program and ancestral-state reconstruction in termites (top), together with a proposed model for the developmental origin of totipotent workers in termites, which evolved from an ancestral condition with linear nymphal development through retention of a juvenile-like transcriptional state (bottom).

Juvenile hormone (JH) and ecdysone signalling, the highly conserved endocrine pathways regulating insect development and differentiation, were also prominent among the ancestral caste DEGs (**Figure 2A**). Allatotropins (*AT*), which encode neuropeptides that can stimulate JH synthesis (Kataoka et al. 1989), JH acid methyltransferase (*jhamt*), which catalyses the final steps of JH biosynthesis, and Krüppel homolog 1 (*Kr-h1*), a highly conserved JH-induced transcription factor (Jindra, Palli, and Riddiford 2013; Belles 2020), as well as *chinmo*, which coordinates with JH to maintain juvenile state in insects (Escudero et al. 2025), were more highly expressed in totipotent workers than nymphs, whereas Juvenile hormone esterase (*Jhe*), which encodes a JH-degrading enzyme, was nymph-biased. These expression patterns indicate higher JH activity in totipotent workers. By contrast, Ecdysone-induced protein 93F (*Eip93F*), a late ecdysone-response transcription factor that specifies the adult developmental program (Ureña et al. 2014; Truman 2019; Belles 2020), showed the strongest nymph- biased expression among the ancestral caste DEGs. Other ecdysteroid-synthesis or ecdysone-responsive genes, including phantom (*phtm*), spook (*spo*), Blimp-1 and ecdysone-inducible gene E1 (*ImpE1*), were also up-regulated in nymphs. Consistent with this pattern, analyses of published termite transcriptomes showed that *Kr-h1* is consistently highly expressed in totipotent workers, whereas *Eip93F* is expressed at comparatively low levels (**Supplementary Figure 4**). These results indicate that high JH activity, coupled with reduced ecdysone-associated metamorphic signalling, is associated with the maintenance of the ancestral totipotent worker state in termites.

Key components and regulators of the TGF-β superfamily signalling pathway, including *myoglianin* (*myo*), *Activin-β*(*Actbeta*), *Sno oncogene* (*Snoo*), and *Follistatin* (*Fs*) (Upadhyay et al. 2017), were also prominent among the nymph-biased ancestral caste DEGs (**Figure 2A**). *Myo* is particularly relevant because it is required for wing-bud formation (Kawamoto et al. 2025) and can suppress JH biosynthesis and regulate ecdysone production in hemimetabolous insects (Ishimaru et al. 2016, 2026). To test the role of TGF-β signalling in termite nymphal differentiation, we collected nymph-developing *Cryptotermes declivis* colonies from the field and used RNA interference to silence *myo* expression. Compared with controls, *myo* knock-down nymphs showed elevated expression level of *Kr-h1*, reduced wing-bud development and delayed metamorphic progression (**Figure 2B**; **Supplementary Figure 5**), indicating that the lower TGF-β superfamily signalling contributes to the high JH level and the maintenance of totipotency in totipotent workers. Together, these results indicate that the evolution of linear caste development in termites involved the co-option of a conserved hemimetabolous insect developmental GRN.

The high JH signalling and low *Eip93F* expression observed in totipotent workers suggest that they are in a juvenile-like developmental state (Grassi and Sandias 1896; C. H. Kennedy 1947; Nalepa and Bandi 2000; Judith Korb and Hartfelder 2008; Nalepa 2026). To test this hypothesis and investigate the developmental origin of totipotent workers in termites, we compared developmental transcriptomes from termites with linear caste development with those from *B. germanica*, a solitary cockroach that undergoes typical nymphal development into winged adults. A PCA of *B. germanica* developmental transcriptomes based on the top 250 termite ancestral caste DEGs revealed a clear nymphal developmental trajectory, with PC1 separating early third-instar nymphs from later instars and PC2 distinguishing the penultimate fifth- from the last (ultimate) sixth-instar nymphs (Figure 2C). This suggests that linear caste development in termites and nymphal development in *B. germanica* involve a shared developmental GRN.

We next projected termite transcriptomes onto this *B. germanica*-trained PCA space (Methods). Totipotent workers mapped near early-instar *B. germanica* nymphs, which primarily undergo feeding and growth (Belles, Maestro, and Piulachs 2024), whereas termite nymphs partially overlapped with totipotent workers and extended toward the penultimate and ultimate *B. germanica* instars (**Figure 2C**), which progressively acquire adult morphology before the final metamorphic moult (Belles, Maestro, and Piulachs 2024). Consistent with this developmental correspondence, *Kr- h1* and *Eip93F* showed sharp expression changes between the penultimate and ultimate nymphal instars of *B. germanica* and between totipotent workers and nymphs across termites, with particularly pronounced shift between early and late nymphal instars of *C. secundus* (**Figure 2D**). In addition, *myo* showed elevated expression in the penultimate and ultimate *B. germanica* instars and was already highly expressed in the early nymphal instar of *C. secundus*, consistent with its upstream role in initiating this developmental transition. In *P. canalifrons* and *Z. nevadensis*, which have only one and two nymphal instars, respectively, these genes also showed greater expression variation among nymphs than among totipotent workers (**Figure 2D**), consistent with endocrine and developmental transitions occurring over a compressed developmental interval. Together, these patterns support the hypothesis that the termite totipotent workers evolved through retention of a juvenile-like developmental state, whereas termite nymphs undergo endocrine activation of the adult developmental program (**Figure 2E**).

### Bifurcated caste development in termites convergently evolved via heterochronic expression shift of the linear caste developmental GRN

In contrast to linear caste development, bifurcated caste development involves early divergence of workers and nymphs into distinct developmental trajectories (Roisin 2000; Judith Korb and Hartfelder 2008). To investigate the gene regulatory basis of these convergent transitions, we identified caste DEGs in species with bifurcated caste development and recovered 2,625 one-to-one orthologs with caste differential expression in at least one species. Consistent with convergent evolution, most genes were species-specific caste DEGs or showed opposite caste-bias directions (**Supplementary Table 3**). However, 148 genes had similar caste-biased expression across all species with bifurcated caste development (hereafter bifurcated-caste DEGs). Many nymph-biased bifurcated-caste DEGs encoded key developmental regulators, including *hedgehog* (*hh*), *engrailed* (*en*), and s*ex combs reduced* (*Scr*), whereas worker-biased bifurcated-caste DEGs were associated with feeding behaviour and synaptic transmission, including *short neuropeptide F* (*sNPF*), *diuretic hormone 31* (*Dh31*), and *orb2* (**Figure 3A**). This divergent expression pattern suggests that the convergent evolution of bifurcated caste development was accompanied with increasing caste specialization, with nymphs biased toward morphogenetic processes associated with their future reproductive role and workers toward colony maintenance functions. This also shows that caste specialization aligns closely with reduced reproductive potential in workers.

**Figure 3.**
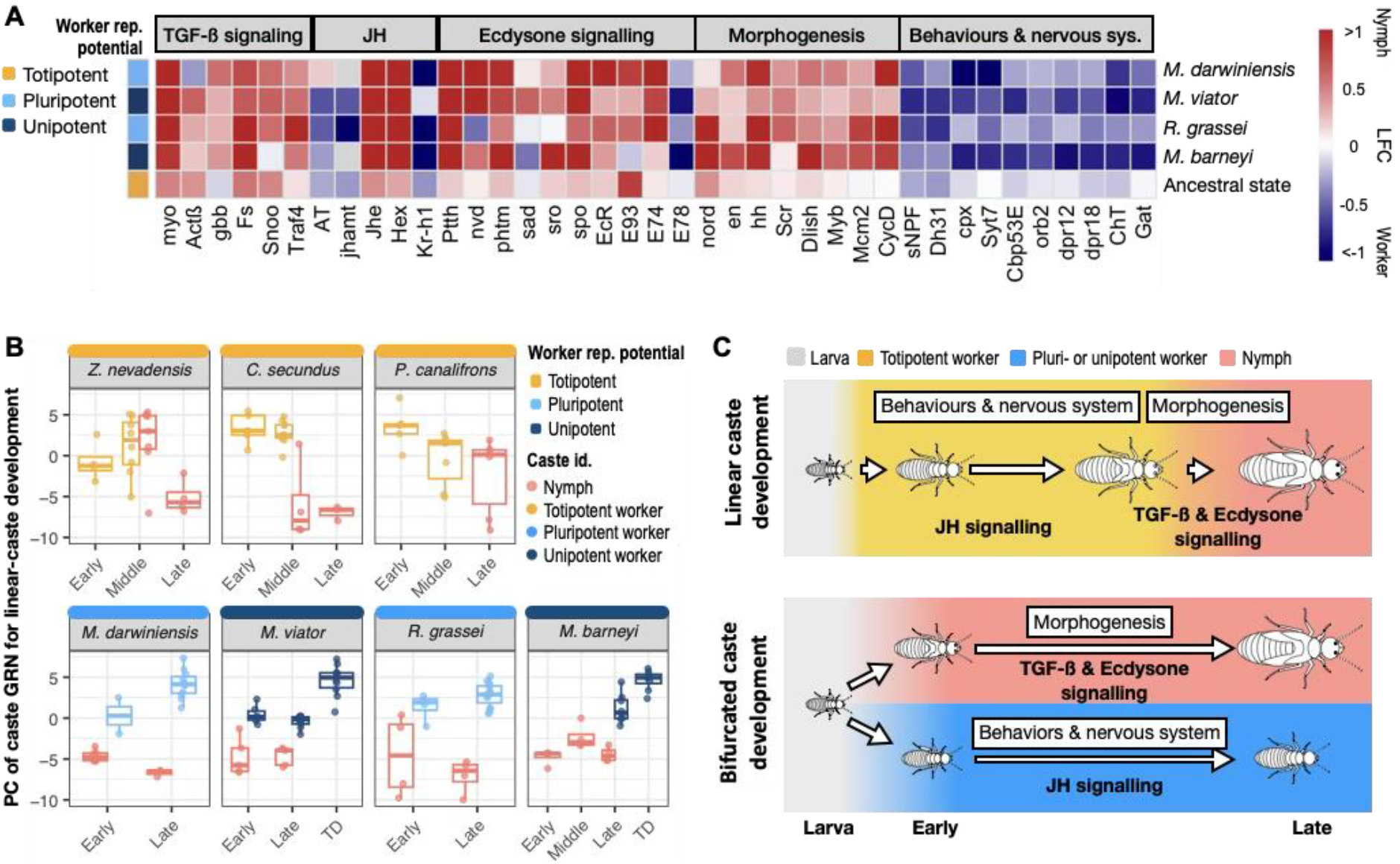
Bifurcated caste development in termites evolved convergently via heterochronic expression shifts. (A) Heatmap showing caste-biased expression of bifurcated-caste DEGs involved in TGF-â, JH, and ecdysone signalling, morphogenesis, and behaviours and nervous system. The reconstructed ancestral caste-biased expression for linear caste development is shown in the bottom row. Additional key regulators of TGF-β and ecdysone signalling are included for comparison. The complete gene list is provided in **Supplementary Table 3**. Colours indicate log2 fold changes (LFC) in expression between castes: red, nymph-biased expression; blue, worker-biased expression; and grey, low expression in both castes (transcripts per million < 2). (B) PC scores representing caste GRN in termites with linear caste development (top row) and its deployment during bifurcated caste development (bottom row). The PC space was trained on developmental transcriptomes of *C. secundus* using the bifurcated-caste DEGs. Corresponding analyses using PC spaces trained on *Z. nevadensis* and *P. canalifrons* are shown in **Supplementary Figure 6**. (C) Proposed model for the developmental origin of bifurcated caste development. Bifurcated caste development evolved repeatedly from linear caste development through recruitment of the ancestral developmental GRN and heterochronic shifts in expression from later to earlier instars.

Key endocrine genes and TGF-β signalling components were represented among the bifurcated caste DEGs (**Figure 3A**). *Jhe* and a hexamerin gene hypothesized to sequester JH (Zhou, Tarver, and Scharf 2006) were among the most strongly nymph-biased bifurcated-caste DEGs, consistent with the higher expression of the JH-signalling gene *Kr-h1* in workers. At the same time, nymphs showed a strong ecdysone signal: key ecdysteroid biosynthesis enzymes, including *phantom* (*phtm*) and *spook*, and the upstream activator prothoracicotropic hormone (*ptth*) (Yamanaka, Rewitz, and O’Connor 2013), were more highly expressed in nymphs than in workers across species, as were the ecdysone receptor (*EcR*) and Ecdysone-induced protein 74EF (*Eip74EF*)—a direct transcriptional target of ecdysone signalling (**Figure 3A**). In addition, the TGF-β signalling components *Fs*, *Snoo*, and *myo* were all nymph-biased among the bifurcated-caste DEGs. Notably, *myo* was significantly differentially expressed between early-stage nymphs and workers in *M. viator*, even before the caste-bias expression of *Kr-h1* became significant, supporting the upstream role of TGF-β signalling. These patterns suggest that bifurcated caste development is associated with high JH signalling in workers and elevated TGF-β and ecdysone signalling for nymphs, resembling the expression contrast between totipotent workers and nymphs observed in termites with linear caste development.

To quantitatively test whether the ancestral developmental GRN underlying linear caste development was repeatedly co-opted during the evolution of bifurcated caste development, we compared transcriptomes from termites across both caste developmental programs. PCA based on the bifurcated-caste DEGs separated totipotent workers from nymphs in termites with linear caste development (**Supplementary Figure 6**), indicating that these genes are associated with worker–nymph differentiation in both developmental programs. We then projected developmental transcriptomes from termites with bifurcated caste development onto a PCA space trained on *C. secundus* transcriptomes (Method; see **Supplementary Figure 6** for equivalent analyses using PCA spaces trained on *Z. nevadensis* or *P. canalifrons*). PC1, which separated totipotent workers and nymphs in termites with linear caste development, also separated workers from nymphs across termites with bifurcated caste development, with caste separation increasing from early to later instars (**Figure 3B**). In particular, early-instar pluripotent and unipotent workers had PC values similar to those of totipotent workers, whereas nymphs of termites with bifurcated caste development resembled late-stage nymphs of termites with linear-caste development (**Figure 3C**). Together, these results suggest that bifurcated caste development evolved through repeated heterochronic recruitment of the ancestral developmental GRN for linear caste development, shifting its activity from later developmental stages to earlier instars and thereby promoting early commitment to worker versus nymphal fate (**Figure 3C**).

### Terminal differentiation transforms unipotent workers into colony-level soma

We next investigated the genetic mechanisms underlying the evolution of unipotent workers, a defining feature of termite superorganismality (Bernadou, Kramer, and Korb 2021). These workers undergo terminal differentiation, after which they lose the capacity to further differentiate into other castes, except for the possibility of developing into pre-soldiers — a pathway limited to young workers and rapidly foreclosed with age (Noirot 1969; Noirot and Pasteels 1987). To this end, we focused on *M. viator* and *M. barneyi*, which represent independent origins of unipotent workers in Hodotermitidae and Termitidae, respectively. We compared worker transcriptomes before and after terminal differentiation and further included soldiers and alates (winged reproductives) from *M. barneyi*, two terminal castes that cannot moult anymore. For comparison with non-terminal workers, we included *P. canalifrons* and *R. grassei*, which represent close relatives of Termitidae that retain totipotent and pluripotent workers, respectively (**Figure 1**).

Consistent with terminal differentiation representing a discrete developmental transition in unipotent workers, PCA clearly separated workers before and after terminal differentiation, and more than 1000 and 4000 genes were differentially expressed between these two stages in *M. viator* and *M. barneyi*, respectively (**Supplementary Table 4**). In both species, genes associated with developmental processes, including DNA replication, neuronal development, and cuticle development were more highly expressed in workers before terminal differentiation, whereas genes involved in oxidative phosphorylation (OXPHOS) were more highly expressed in terminally differentiated workers (**Figure 4A**). Notably, parallel expression changes occurred during the imaginal moult into alates and the terminal moult into soldiers: workers before terminal differentiation resembled late-instar nymphs and pre-soldiers, whereas terminally differentiated workers resembled alates and soldiers. In contrast, opposite developmental expression patterns were found in *P. canalifrons* and *R. grassei* (**Figure 4A**), suggesting distinct developmental trajectories between terminal and non-terminal worker castes.

**Figure 4.**
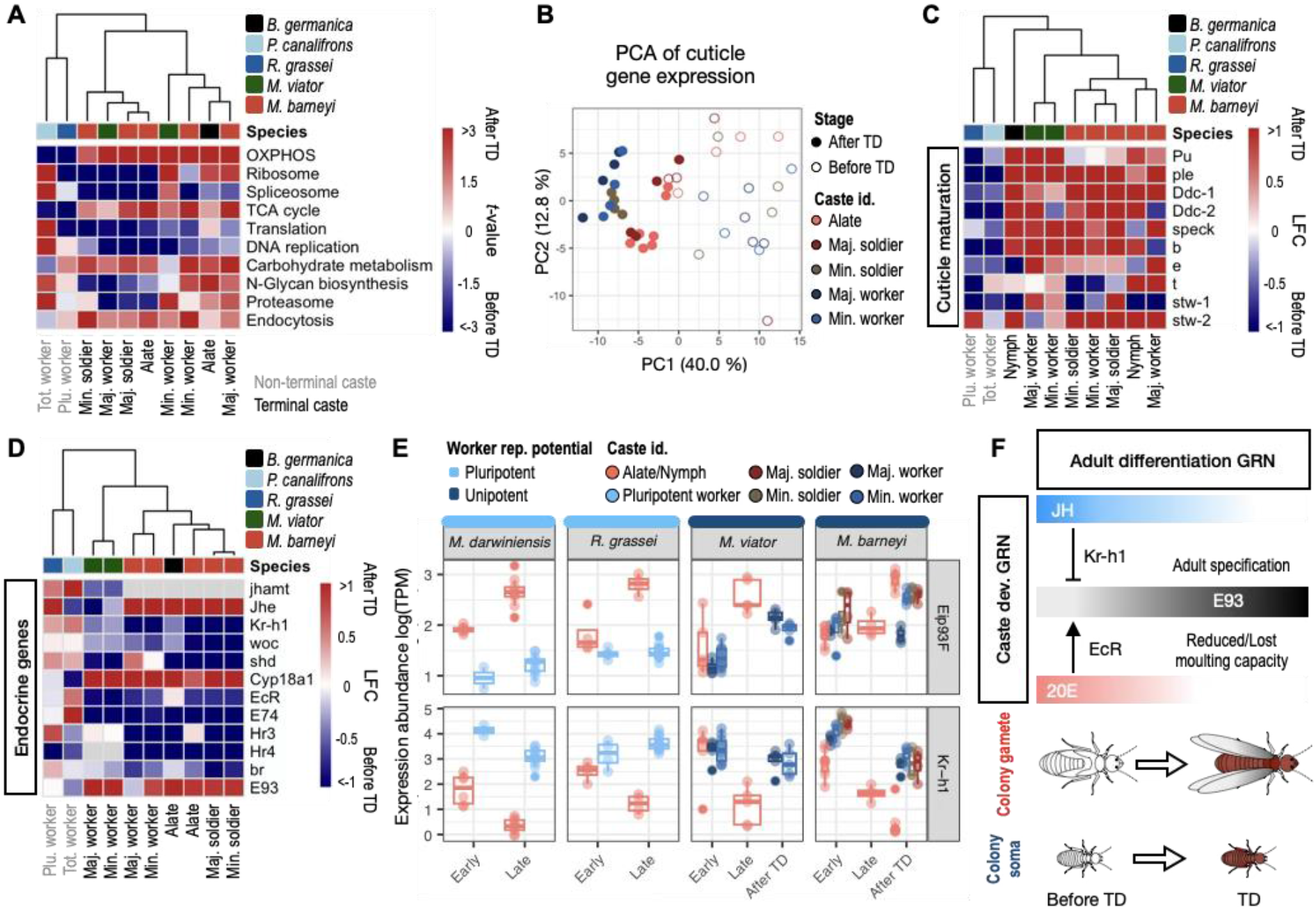
Unipotent workers in termites evolved through co-option of an endocrine GRN for adult differentiation. (A) Heatmap showing KEGG pathway enrichment associated with gene expression changes during terminal differentiation (TD) in termite species with unipotent workers. For comparison, equivalent analyses are shown for *R. grassei* and *P. canalifrons*, which have non-terminal workers, and for *B. germanica*. Colours indicate *t*-values from gene-set enrichment analyses: red, higher expression after TD; blue, higher expression before TD; grey, low expression (TPM < 2). Species are indicated by the top colour bar. For *R. grassei* and *P. canalifrons*, the contrasts are between late- and early-instar workers. (B) PCA of cuticle-related gene expression across workers, soldiers, and nymphs of *M. barneyi*. Colours indicate caste, and symbol fill indicates developmental stage: open symbols, before TD; filled symbols, after TD. A corresponding analysis for *M. viator* workers is shown in **Supplementary Figure 7**. (C) Heatmap showing stage-associated expression differences in genes involved in dopamine-mediated cuticle maturation, including melanisation and sclerotization. For genes duplicated in termites, both copies are shown. (D) Heatmaps showing stage-associated expression differences in genes belonging to the endocrine GRN underlying adult differentiation in insects. (E) Expression of *E93* and *Kr-h1*, shown as log2-transformed TPM, across development or terminal differentiation in workers, soldiers, and nymphs from termite species with pluripotent or unipotent workers. (F) Proposed model for the developmental origin of unipotent workers. Worker terminal differentiation evolved through co-option of the endocrine GRN regulating the nymph-to-adult transition (adult differentiation GRN), with caste-specific morphology being specified by caste-developmental GRNs acting beforehand. For C and D, the contrasts for *R. grassei* and *P. canalifrons* are as in panel A.

A similar pattern was observed for genes involved in cuticle sclerotization and melanisation, processes that are characteristic of unipotent workers, alates, and soldiers but absent or less pronounced in totipotent and pluripotent workers (Noirot 1969). In both *M. viator* and *M. barneyi*, PCA based on the expression of genes encoding cuticle proteins and those involved in cuticle maturation clearly separated workers before and after terminal differentiation (Figure 4B) (**Supplementary Figure 7**), consistent with their distinct cuticular phenotypes (Noirot 1969). In particular, key enzymes involved in dopamine biosynthesis, including *pale*, *punch*, and *dopa decarboxylase* (*Ddc*), were highly expressed in terminally differentiated workers in both species (**Figure 4C**), suggesting increased production of dopamine, the direct biochemical precursor of cuticular melanization and sclerotization in insects (Noh et al. 2016; Sugumaran 2022). Consistently, downstream components of the dopamine-mediated cuticle maturation pathway, such as *speck* and *ebony*, also showed elevated expression in terminally differentiated workers (**Figure 4C**), although the magnitude of these changes differed between the two species, suggesting lineage-specific modulation of the pathway. Interestingly, these genes also exhibited parallel expression changes during terminal differentiation in soldiers and alates, but showed the opposite expression bias during worker development in totipotent and pluripotent workers (**Figure 4C**). Because elevated OXPHOS-related expression is consistent with an adult-like metabolic state (Wilkens et al. 2025), and strong cuticular sclerotization and melanisation are hallmarks of insect adult differentiation (Sugumaran 2022), these findings indicate that terminal differentiation in unipotent workers involves a coordinated shift toward an adult-like transcriptomic state. This contrasts with totipotent workers and pluripotent workers, which remain in a juvenile-like state, and it marks the irreversible transformation of unipotent workers into colony soma.

### Endocrine GRN co-option underlies terminal differentiation of unipotent workers

The parallel transcriptomic shifts observed across terminal castes led us to hypothesize that terminal differentiation in unipotent workers evolved through co-option of an endocrine GRN that normally regulates the nymph-to-adult transition in hemimetabolous insects. To test this hypothesis, we examined the expression of core endocrine genes underlying adult differentiation in insects. Although their overall expression profiles differed among castes (**Supplementary Figure 8**), multiple components showed parallel changes during terminal differentiation in unipotent workers, soldiers, and alates (**Figure 4D**). *EcR*, *Eip74EF*, and *without children* (*woc*), which encodes a transcription factor that controls ecdysone biosynthesis (Lafont et al. 2012), were consistently more highly expressed before terminal differentiation. By contrast, *Cyp18a1*, which encodes a 20-Hydroxyecdysone (20E)- inactivating enzyme, was upregulated afterward. These patterns suggest activation of the ecdysone-responsive moulting program before terminal differentiation, followed by 20E clearance after the terminal moult, in line with the strongly reduced moulting capacity and rudimentary prothoracic glands of terminally differentiated unipotent workers (Okot-Kotber 1980). Terminal differentiation in unipotent workers was also accompanied by reduced *Kr-h1*, indicating decreased JH signalling, consistent with the reduced corpora allata of terminally differentiated workers in *Macrotermes* (Okot-Kotber 1980). This endocrine shift resembled that associated with the imaginal moult in alates and the terminal moult in soldiers (**Figure 4E**) (Okot-Kotber 1985). However, before terminal differentiation, unipotent workers and soldiers maintained higher *Kr-h1* expression than nymphs (**Figure 4E**), suggesting that this endocrine GRN is deployed within caste-specific developmental contexts (**Figure 4F**).

*Eip93F* was among the genes most strongly associated with terminal differentiation and was upregulated more than twofold in unipotent workers of both *M. viator* and *M. barneyi*, except in *M. barneyi* major workers, potentially reflecting their capacity to undergo limited stationary moults (Noirot 1969). This pattern mirrored the high *Eip93F* expression observed in soldiers, alates, and adults *B. germanica*, but contrasted with its persistently low expression in totipotent and pluripotent workers (**Figure 4E**). Because Eip93F is an adult-specifier (Ureña et al. 2014), these patterns suggest that its upregulation contributes to the adult-like transcriptomic state of terminally differentiated unipotent workers. In insects, *Eip93F* is induced by ecdysone signalling and repressed by JH-induced *Kr-h1* (Kayukawa et al. 2017). The concurrent increase in *Eip93F* expression, activation of the ecdysone-response pathway, and decline of JH signalling therefore indicate deployment of the conserved endocrine switch controlling adult differentiation. Consistent with this regulatory relationship, we identified Kr-h1-binding site like sequences in the Eip93F promoters across all termite species examined (**Method**) (**Supplementary Table 5**). Together, these findings indicate that the evolution of unipotent workers involved co-option of a conserved endocrine GRN that underlies adult differentiation in insects (**Figure 4F**).

## Discussion

### The evolutionary history of METs to superorganismal termites

For more than 40 years the ancestral worker state in termites is debated (Watson and Sewell 1985; Noirot and Pasteels 1987; Thompson et al. 2004; Grandcolas and D’Haese 2004; Legendre et al. 2008; Hellemans et al. 2024), largely prompted by the detection of bifurcated caste development in the phylogenetically most basal extant termite *M. darwiniensis* (Watson and Sewell 1985). By integrating comparative developmental genetics with targeted sampling of key termite lineages spanning different degrees of workers’ reproductive potential and several transition nodes (**Figure 1**), we reconstructed, for the first time, the evolutionary trajectory leading to superorganisms in termites. Ancestral-state reconstruction indicates that linear caste development represents the ancestral developmental mode in termites, consistent with the most recent phylogenetic and taxonomic revision of the group (Hellemans et al. 2024). We identified at least three independent origins: in the lineage leading to (i) Mastotermitidae, where totipotent workers evolved into pluripotent workers; (ii) Hodotermitidae, where totipotent workers likely evolved into unipotent workers via an intermediate pluripotent stage in the unsampled *Anacanthotermes* lineage; and (iii) Heterotermitidae, where totipotent workers evolved into pluripotent workers, followed by the evolution of sterile, unipotent workers in the ancestor of Termitidae.

Our inferred evolutionary transition from toti- to pluri- and unipotent workers indicates gradual loss of workers’ reproductive potential. This is associated with increasing evolutionary reproductive altruism, increasing importance of indirect compared to direct fitness benefits, and decreasing (potential) conflict within colonies (Lohmar et al. 2026). Analogously, also at the transition from unicellular eukaryotes to multicellular animals the degree of reproductive altruism of lower-level altruistic units, somatic cells, increased. Somatic cells lose potency from toti- to pluri- and unipotent cells through increasingly developmentally restricted somatic cell types (Galis, Metz, and Alphen 2018). In multicellular animals, these trajectories are accompanied by germline-soma segregation occurring at increasingly earlier stages during development (leading to the Weismann barrier), thereby restricting the developmental potential of somatic cells and enabling their functional specialization (Buss 1987; C. G. Extavour and Akam 2003; C. G. M. Extavour 2007). Likewise, our results indicate that the evolution of termite superorganismality involved an analogous earlier commitment of individuals to the worker versus reproductive developmental pathways and an increasing developmental canalization and functional transcriptomic specialization between castes. These striking similarities across both METs point to early determination of fate as a key mechanism that proximately enables specialization and which ultimately also reduces conflicts.

### Mechanisms associated with the transitions to superorganimality

The reduction of workers’ reproductive potential during the METs to superorganismality in termites was achieved through recurrent co-option of ancestral developmental GRNs coupled with heterochronic shifts in their deployment.

Our ancestral GRN reconstruction indicates that the linear caste development of termites with totipotent workers originated through co-option of an ancestral nymphal developmental GRN from solitary cockroaches, analogous to the co-option of pre-exiting key transcription factors during the evolution of simple multicellularity from unicellular ancestors (Mendoza et al. 2013). This conserved nymphal developmental GRN comprises wing and eye developmental genes (e.g. *vg*, *ap*, *bi*), key components and regulators of the TGF-β superfamily signalling pathway (e.g. *Actbeta*, *Snoo*, *myo*, *Fs*), and an associated endocrine signal, typical of nymphal development in cockroaches. In addition, we showed that, consistent with a conserved nymphal developmental GRN, TGF-β signalling, in particular myo, plays an upstream regulatory role for the onset of nymphal development in linear caste species. These findings suggest that a low TGF-β signalling maintains worker totipotency by sustaining high JH titres and suppressing ecdysone signalling. This is supported by endocrine data showing that nymphal development is associated with decreasing JH titres (Cornette et al. 2008; Judith Korb, Hoffmann, and Hartfelder 2009, 2012). Overall, the timing of the gene expression implies that (wingless) totipotent workers represent an extension of cockroaches’ early developmental stages with delayed onset of morphogenesis. This supports earlier hypotheses proposing that, during the evolution of termites from wood-nesting cockroach ancestors, developmental timing and ontogenetic patterning were modified, with termite workers corresponding to extended early cockroach stages (Judith Korb, Hoffmann, and Hartfelder 2009; Nalepa 2010).

During subsequent stages of the METs in termites, bifurcated caste development evolved repeatedly through recurrent co-option of the same developmental GRN via heterochronic shifts in expression to earlier developmental stages. This heterochronic shift partitioned a previously linear developmental trajectory into alternative developmental pathways, in which individuals either follow the nymphal path towards dispersing reproductives or the wingless worker trajectory. This heterochronic shift forms the mechanistic basis earlier caste fate commitment, associated with increased specialization and conflict reduction. It is analogous to the partitioning of temporally alternating phenotypes in unicellular organisms into spatially differentiated cell types during the evolution of multicellularity (Brunet and King 2017). A comparable bifurcation of developmental trajectories evolved independently in superorganismal social Hymenoptera. where larvae likewise commit to either reproductive or worker developmental pathways. The latter are characterized by a suppression of reproductive organ development and in ants, in addition, by inhibited wing development (Wheeler 1986). However, the underlying regulatory mechanisms differ fundamentally between holometabolous Hymenoptera and hemimetabolous termites. While termite workers retain high JH signalling (Cui et al. 2026), worker development in ants and honey bees is associated with reduced JH signalling (Hartfelder 2000; Li et al. 2024; Kocher and Kingwell 2024; Shalem, Goldberg, and Bloch 2025), probably reflecting the distinct developmental architectures of hemimetabolous and holometabolous insects. Thus, although major transitions repeatedly converge on reduced workers’ reproductive potential, the developmental mechanisms that achieve this convergence remain strongly contingent on lineage-specific ancestral developmental programs.

The transitions to superorganismal termites—the independent evolution of sterile, unipotent workers in Termitidae and Hodotermitidae—was achieved through terminal differentiation of workers. In contrast to totipotent and pluripotent workers, which retain a juvenile-like state and maintain full or partial reproductive potential, respectively, our results indicate that unipotent workers adopt an adult-like developmental state. This is evidenced by strong oxidative phosphorylation and cuticle maturation signatures, shared with terminally differentiated soldiers and alates, and workers in the holometabolous social Hymenoptera. This evolutionary innovation in unipotent termite workers was mediated by repeated co-option of the endocrine GRN governing adult differentiation in hemimetabolous insects, which is characterized by the adult-specifier transcription factor Eip93F. Worker terminal differentiation irreversibly eliminates developmental plasticity and reproductive potential and is accompanied by functional specialization, analogous to the evolution of terminally differentiated somatic cells in metazoan multicellularity. Thus, it provides the mechanistic basis for resolving reproductive conflict and elaborating division of labour, which may form a mutually reinforcing cycle, transforming colonies into superorganisms.

## References

Belles, Xavier. 2020. ‘Krüppel Homolog 1 and E93: The Doorkeeper and the Key to Insect Metamorphosis’. Archives of Insect Biochemistry and Physiology 103 (3): e21609. 10.1002/arch.21609.

Belles, Xavier, Jose Luis Maestro, and Maria-Dolors Piulachs. 2024. ‘The German Cockroach as a Model in Insect Development and Reproduction in an Endocrine Context’. Advances in Insect Physiology, 2024, 1–47. 10.1016/bs.aiip.2024.03.001.

Berens, Ali J., James H. Hunt, and Amy L. Toth. 2014. ‘Comparative Transcriptomics of Convergent Evolution: Different Genes but Conserved Pathways Underlie Caste Phenotypes across Lineages of Eusocial Insects’. Molecular Biology and Evolution 32 (3): 690–703. 10.1093/molbev/msu330.

Bernadou, Abel, Boris H. Kramer, and Judith Korb. 2021. ‘Major Evolutionary Transitions in Social Insects, the Importance of Worker Sterility and Life History Trade-Offs’. Frontiers in Ecology and Evolution 9 (2021): 732907. 10.3389/fevo.2021.732907.

Boomsma, Jacobus J., and Richard Gawne. 2018. ‘Superorganismality and Caste Differentiation as Points of No Return: How the Major Evolutionary Transitions Were Lost in Translation’. Biological Reviews 93 (1): 28–54. 10.1111/brv.12330.

Brawand, David, Magali Soumillon, Anamaria Necsulea, Philippe Julien, Gábor Csárdi, Patrick Harrigan, Manuela Weier, et al. 2011. ‘The Evolution of Gene Expression Levels in Mammalian Organs’. Nature 478 (7369): 343–48. 10.1038/nature10532.

Brunet, Thibaut, and Nicole King. 2017. ‘The Origin of Animal Multicellularity and Cell Differentiation.’ Developmental Cell 43 (2): 124–40. 10.1016/j.devcel.2017.09.016.

Buss, Leo W. 1983. ‘Evolution, Development, and the Units of Selection.’ Proceedings of the National Academy of Sciences 80 (5): 1387–91. 10.1073/pnas.80.5.1387.

Buss, Leo W. 1987. The Evolution of Individuality. Princeton University Press. JSTOR. http://www.jstor.org/stable/j.ctt7zvwtj.

Cornette, Richard, Hiroki Gotoh, Shigeyuki Koshikawa, and Toru Miura. 2008. ‘Juvenile Hormone Titers and Caste Differentiation in the Damp-Wood Termite Hodotermopsis Sjostedti (Isoptera, Termopsidae)’. Journal of Insect Physiology 54 (6): 922–30. 10.1016/j.jinsphys.2008.04.017.

Cui, Yingying, Fangfang Liu, Dongwei Yuan, Mingtao Liao, Zhaoxin Li, Yun-Xia Luan, Shuxin Yu, et al. 2026. ‘Nutritional Specialization and Social Evolution in Woodroaches and Termites’. Science (New York, N.Y.) 392 (6794): eadt2178. 10.1126/science.adt2178.

Devlin, Dominic K., Austen R. D. Ganley, and Nobuto Takeuchi. 2023. ‘A Pan-Metazoan View of Germline-Soma Distinction Challenges Our Understanding of How the Metazoan Germline Evolves’. Current Opinion in Systems Biology 36 (2023): 100486. 10.1016/j.coisb.2023.100486.

Escudero, Jorge, Judit Gonzalvo, Maria-Dolors Piulachs, and Xavier Belles. 2025. ‘Chinmo Function in Cockroaches Provides New Insights into the Regulation and Evolution of Insect Metamorphosis’. PLOS Genetics 21 (12): e1011993. 10.1371/journal.pgen.1011993.

Evans, Jay D., and Diana E. Wheeler. 1999. ‘Differential Gene Expression between Developing Queens and Workers in the Honey Bee, Apis Mellifera’. Proceedings of the National Academy of Sciences 96 (10): 5575–80. 10.1073/pnas.96.10.5575.

Extavour, Cassandra G., and Michael Akam. 2003. ‘Mechanisms of Germ Cell Specification across the Metazoans: Epigenesis and Preformation’. Development 130 (24): 5869–84. 10.1242/dev.00804.

Extavour, Cassandra G. M. 2007. ‘Evolution of the Bilaterian Germ Line: Lineage Origin and Modulation of Specification Mechanisms’. Integrative and Comparative Biology 47 (5): 770–85. 10.1093/icb/icm027.

Feigin, Charles, Sha Li, Jorge Moreno, and Ricardo Mallarino. 2023. ‘The GRN Concept as a Guide for Evolutionary Developmental Biology’. Journal of Experimental Zoology Part B: Molecular and Developmental Evolution 340 (2): 92–104. 10.1002/jez.b.23132.

Galis, Frietson, Johan A. J. Metz, and Jacques J. M. van Alphen. 2018. ‘Development and Evolutionary Constraints in Animals’. *Annual Review of Ecology*, Evolution, and Systematics 49 (1): 499–522. 10.1146/annurev-ecolsys-110617-062339.

Gardner, A., and A. Grafen. 2009. ‘Capturing the Superorganism: A Formal Theory of Group Adaptation’. Journal of Evolutionary Biology 22 (4): 659–71. 10.1111/j.1420-9101.2008.01681.x.

Grandcolas, P., and C. D’Haese. 2004. ‘The Origin of a “True” Worker Caste in Termites: Mapping the Real World on the Phylogenetic Tree’. Journal of Evolutionary Biology 17 (2): 461–63. 10.1046/j.1420-9101.2003.00662.x.

Grassi, B., and A. Sandias. 1896. The Constitution and Development of the Society Of Termites: Observations On Their Habits; With Appendices on the Parasitic Protozoa Of Termitidæ, and on The Embiidæ. s2-39 (155): 245–322. 10.1242/jcs.s2-39.155.245.

Hamilton, W. D. 1964a. ‘The Genetical Evolution of Social Behaviour. I’. Journal of Theoretical Biology 7 (1): 1–16. 10.1016/0022-5193(64)90038-4.

Hamilton, W. D. 1964b. ‘The Genetical Evolution of Social Behaviour. II’. Journal of Theoretical Biology 7 (1): 17–52. 10.1016/0022-5193(64)90039-6.

Hartfelder, K. 2000. ‘Insect Juvenile Hormone: From “Status Quo” to High Society’. Brazilian Journal of Medical and Biological Research 33 (2): 157–77. 10.1590/s0100-879x2000000200003.

Hellemans, Simon, Mauricio M. Rocha, Menglin Wang, Johanna Romero Arias, Duur K. Aanen, Anne-Geneviève Bagnères, Aleš Buček, et al. 2024. ‘Genomic Data Provide Insights into the Classification of Extant Termites’. Nature Communications 15 (1): 6724. 10.1038/s41467-024-51028-y.

Howe, Jack, Jochen C. Rink, Bo Wang, and Ashleigh S. Griffin. 2022. ‘Multicellularity in Animals: The Potential for within-Organism Conflict’. Proceedings of the National Academy of Sciences 119 (32): e2120457119. 10.1073/pnas.2120457119.

Ishimaru, Yoshiyasu, Kohei Kawamoto, Sumihare Noji, and Taro Mito. 2026. ‘TGF-β-Dependent Regulation of Juvenile Hormone Biosynthesis in Insect Development and Metamorphosis’. Current Opinion in Insect Science 75 (2026): 101490. 10.1016/j.cois.2026.101490.

Ishimaru, Yoshiyasu, Sayuri Tomonari, Yuji Matsuoka, Takahito Watanabe, Katsuyuki Miyawaki, Tetsuya Bando, Kenji Tomioka, Hideyo Ohuchi, Sumihare Noji, and Taro Mito. 2016. ‘TGF-β Signaling in Insects Regulates Metamorphosis via Juvenile Hormone Biosynthesis’. Proceedings of the National Academy of Sciences 113 (20): 5634–39. 10.1073/pnas.1600612113.

Jindra, Marek, Subba R. Palli, and Lynn M. Riddiford. 2013. ‘The Juvenile Hormone Signaling Pathway in Insect Development’. Annual Review of Entomology 58 (1): 181–204. 10.1146/annurev-ento-120811-153700.

Kataoka, Hiroshi, Anne Toschi, Jorge P. Li, Robert L. Carney, David A. Schooley, and Steven J. Kramer. 1989. ‘Identification of an Allatotropin from Adult Manduca Sexta’. Science 243 (4897): 1481–83. 10.1126/science.243.4897.1481.

Kawamoto, Kohei, Yoshiyasu Ishimaru, Sayuri Tomonari, Takahito Watanabe, Sumihare Noji, and Taro Mito. 2025. ‘Myoglianin Is a Crucial Factor for the Transition to the Juvenile Hormone-Dependent Phase during Hemimetabolous Nymphal Development’. Insect Biochemistry and Molecular Biology 178 (2025): 104274. 10.1016/j.ibmb.2025.104274.

Kayukawa, Takumi, Akiya Jouraku, Yuka Ito, and Tetsuro Shinoda. 2017. ‘Molecular Mechanism Underlying Juvenile Hormone-Mediated Repression of Precocious Larval–Adult Metamorphosis’. Proceedings of the National Academy of Sciences 114 (5): 1057–62. 10.1073/pnas.1615423114.

Kennedy, Clarence Hamilton. 1947. ‘Child Labor of the Termite Society versus Adult Labor of the Ant Society’. The Scientific Monthly 65 (4): 309. JSTOR. http://www.jstor.org/stable/19227.

Kennedy, Patrick, Gemma Baron, Bitao Qiu, Dalial Freitak, Heikki Helanterä, Edmund R. Hunt, Fabio Manfredini, et al. 2017. ‘Deconstructing Superorganisms and Societies to Address Big Questions in Biology’. Trends in Ecology & Evolution 32 (11): 861–72. 10.1016/j.tree.2017.08.004.

Kocher, Sarah, and Callum Kingwell. 2024. ‘The Molecular Substrates of Insect Eusociality’. Annual Review of Genetics 58 (1): 273–95. 10.1146/annurev-genet-111523-102510.

Korb, J. 2025. ‘Changes of Division of Labour along the Eusociality Spectrum in Termites, with Comparisons to Multicellularity’. Philosophical Transactions B 380 (1922): 20230268. 10.1098/rstb.2023.0268.

Korb, Judith. 2025. ‘Cooperation and Conflict in Termite Societies.’ Current Opinion in Insect Science 71 (2025): 101401. 10.1016/j.cois.2025.101401.

Korb, Judith, and Klaus Hartfelder. 2008. ‘Life History and Development - a Framework for Understanding Developmental Plasticity in Lower Termites’. Biological Reviews 83 (3): 295–313. 10.1111/j.1469-185x.2008.00044.x.

Korb, Judith, Katharina Hoffmann, and Klaus Hartfelder. 2009. ‘Endocrine Signatures Underlying Plasticity in Postembryonic Development of a Lower Termite, Cryptotermes Secundus (Kalotermitidae)’. Evolution & Development 11 (3): 269–77. 10.1111/j.1525-142x.2009.00329.x.

Korb, Judith, Katharina Hoffmann, and Klaus Hartfelder 2012. ‘Molting Dynamics and Juvenile Hormone Titer Profiles in the Nymphal Stages of a Lower Termite, Cryptotermes Secundus (Kalotermitidae) – Signatures of Developmental Plasticity’. Journal of Insect Physiology 58 (3): 376–83. 10.1016/j.jinsphys.2011.12.016.

Lafont, Rene, C. Dauphin-Villemant, J. T. Warren, and H. Rees. 2012. 4 - Ecdysteroid Chemistry and Biochemistry. Edited by [“Lawrence I. Gilbert”]. San Diego: Academic Press. 10.1016/b978-0-12-384749-2.10004-4.

Legendre, Frédéric, Michael F. Whiting, Christian Bordereau, Eliana M. Cancello, Theodore A. Evans, and Philippe Grandcolas. 2008. ‘The Phylogeny of Termites (Dictyoptera: Isoptera) Based on Mitochondrial and Nuclear Markers: Implications for the Evolution of the Worker and Pseudergate Castes, and Foraging Behaviors’. Molecular Phylogenetics and Evolution 48 (2): 615–27. 10.1016/j.ympev.2008.04.017.

Li, Ruyan, Xueqin Dai, Jixuan Zheng, Rasmus Stenbak Larsen, Yanmei Qi, Xiafang Zhang, Joel Vizueta, Jacobus J. Boomsma, Weiwei Liu, and Guojie Zhang. 2024. ‘Juvenile Hormone as a Key Regulator for Asymmetric Caste Differentiation in Ants’. Proceedings of the National Academy of Sciences 121 (46): e2406999121. 10.1073/pnas.2406999121.

Lohmar, Stephan, Julius Rombach, Louis Allan Okwaro, Wei Zhou, Volker Nehring, and Judith Korb. 2026. ‘A Unifying Framework for the Evolution of Metazoa and Social Insects, Two Major Evolutionary Transitions’. Proceedings of the National Academy of Sciences 123 (32): e2530125123. 10.1073/pnas.2530125123.

Mendoza, Alex de, Arnau Sebé-Pedrós, Martin Sebastijan Šestak, Marija Matejčić, Guifré Torruella, Tomislav Domazet-Lošo, and Iñaki Ruiz-Trillo. 2013. ‘Transcription Factor Evolution in Eukaryotes and the Assembly of the Regulatory Toolkit in Multicellular Lineages’. Proceedings of the National Academy of Sciences 110 (50): E4858–66. 10.1073/pnas.1311818110.

Nalepa, Christine A. 2010. ‘Altricial Development in Subsocial Cockroach Ancestors: Foundation for the Evolution of Phenotypic Plasticity in Termites’. Evolution & Development 12 (1): 95–105. 10.1111/j.1525-142x.2009.00394.x.

Nalepa, Christine A. 2026. ‘Origins of Termite Eusociality: Developmental Foundations’. Frontiers in Ecology and Evolution 14 (2026): 1836415. 10.3389/fevo.2026.1836415.

Nalepa, Christine A., and Claudio Bandi. 2000. Characterizing the Ancestors: Paedomorphosis and Termite Evolution. Dordrecht: Springer Netherlands. 10.1007/978-94-017-3223-9_3.

Noh, Mi Young, Subbaratnam Muthukrishnan, Karl J. Kramer, and Yasuyuki Arakane. 2016. ‘Cuticle Formation and Pigmentation in Beetles’. Current Opinion in Insect Science 17 (2016): 1–9. 10.1016/j.cois.2016.05.004.

Noirot, C. 1969. 10 - Formation of Castes in the Higher Termites*\*\**Translated from the French by Mina Parsont and Frances M. Weesner. Edited by [“Kumar Krishna” and “Frances M. Weesner”]. Academic Press. 10.1016/b978-0-12-395529-6.50014-3.

Noirot, C., and J. M. Pasteels. 1987. ‘Ontogenetic Development and Evolution of the Worker Caste in Termites’. Experientia 43 (8): 851–60. 10.1007/bf01951642.

Okot-Kotber, B. M. 1980. ‘Histological and Size Changes in Corpora Allata and Prothoracic Glands during Development ofMacrotermes Michaelseni (Isoptera)’. Insectes Sociaux 27 (4): 361–76. 10.1007/bf02223729.

Okot-Kotber, B. M. 1985. CHAPTER 21 - Mechanisms of Caste Determination in a Higher Termite, Macrotermes Michaelseni (Isoptera, Macrotermitinae). Edited by [“J.A.L. Watson”, “B.M. Okot- Kotber”, and “CH. Noirot”]. Amsterdam: Pergamon. 10.1016/b978-0-08-030783-1.50026-4.

Qiu, Bitao, Xueqin Dai, Panyi Li, Rasmus Stenbak Larsen, Ruyan Li, Alivia Lee Price, Guo Ding, et al. 2022. ‘Canalized Gene Expression during Development Mediates Caste Differentiation in Ants’. Nature Ecology & Evolution 6 (11): 1753–65. 10.1038/s41559-022-01884-y.

Qiu, Bitao, Rasmus Stenbak Larsen, Ni-Chen Chang, John Wang, Jacobus J. Boomsma, and Guojie Zhang. 2018. ‘Towards Reconstructing the Ancestral Brain Gene-Network Regulating Caste Differentiation in Ants’. Nature Ecology & Evolution 2 (11): 1782–91. 10.1038/s41559-018-0689-x.

Roisin, Yves. 2000. Diversity and Evolution of Caste Patterns. Dordrecht: Springer Netherlands. 10.1007/978-94-017-3223-9_5.

Roisin, Yves, and Judith Korb. 2011. Social Organisation and the Status of Workers in Termites. Dordrecht: Springer Netherlands. 10.1007/978-90-481-3977-4_6.

Shalem, Yuval, Tzvi S. Goldberg, and Guy Bloch. 2025. ‘Juvenile Hormone Signaling and Social Complexity in the Hymenoptera’. Current Opinion in Insect Science 72 (2025): 101433. 10.1016/j.cois.2025.101433.

Sugumaran, Manickam. 2022. ‘Cuticular Sclerotization in Insects – A Critical Review’. Advances in Insect Physiology, 2022, 111–214. 10.1016/bs.aiip.2022.02.001.

Szathmary, Eors, and John Maynard Smith. 1995. ‘The Major Evolutionary Transitions’. Nature 374 (6519): 227--232. 10.1038/374227a0.

Thompson, G. J., O. Kitade, N. Lo, and R. H. Crozier. 2004. ‘On the Origin of Termite Workers: Weighing up the Phylogenetic Evidence’. Journal of Evolutionary Biology 17 (1): 217–20. 10.1046/j.1420-9101.2003.00645.x.

Tripathi, Bipin Kumar, and Kenneth D. Irvine. 2021. ‘The Wing Imaginal Disc.’ Genetics 220 (4): iyac020. 10.1093/genetics/iyac020.

Truman, James W. 2019. ‘The Evolution of Insect Metamorphosis’. Current Biology 29 (23): R1252–68. 10.1016/j.cub.2019.10.009.

Upadhyay, Ambuj, Lindsay Moss-Taylor, Myung-Jun Kim, Arpan C. Ghosh, and Michael B. O’Connor. 2017. ‘TGF-β Family Signaling in Drosophila’. Cold Spring Harbor Perspectives in Biology 9 (9): a022152. 10.1101/cshperspect.a022152.

Ureña, Enric, Cristina Manjón, Xavier Franch-Marro, and David Martín. 2014. ‘Transcription Factor E93 Specifies Adult Metamorphosis in Hemimetabolous and Holometabolous Insects’. Proceedings of the National Academy of Sciences 111 (19): 7024–29. 10.1073/pnas.1401478111.

Wagner, Günter P. 2007. ‘The Developmental Genetics of Homology’. Nature Reviews Genetics 8 (6): nrg2099. 10.1038/nrg2099.

Warner, Michael R., Lijun Qiu, Michael J. Holmes, Alexander S. Mikheyev, and Timothy A. Linksvayer. 2019. ‘Convergent Eusocial Evolution Is Based on a Shared Reproductive Groundplan plus Lineage-Specific Plastic Genes’. Nature Communications 10 (1): 2651. 10.1038/s41467-019-10546-w.

Watson, J. A. L., and J. J. Sewell. 1985. Caste Development in Mastotermes and Kalotermes: Which Is Primitive? Edited by [“J.A.L. Watson”, “B.M. Okot-Kotber”, and “CH. Noirot”]. Section B: Pathways of Caste Development in Principal Termite Groups. Amsterdam: Pergamon. 10.1016/b978-0-08-030783-1.50008-2.

West, Stuart A., Roberta M. Fisher, Andy Gardner, and E. Toby Kiers. 2015. ‘Major Evolutionary Transitions in Individuality’. Proceedings of the National Academy of Sciences 112 (33). 10.1073/pnas.1421402112.

Wheeler, Diana E. 1986. ‘Developmental and Physiological Determinants of Caste in Social Hymenoptera: Evolutionary Implications’. American Society of Naturalists, ahead of print, 1986. 10.1086/284536;issue:issue:10.2307/i320756;pagegroup:string:publication.

Wilkens, Maya, Susanne Zimbelmann, Franziska Roth, Jasmin Cartano, Sergi Sayols, Mario Dejung, Michal Levin, and Falk Butter. 2025. ‘Unraveling Developmental Gene Regulation in Holometabolous Insects through Comparative Transcriptomics and Proteomics’. Communications Biology 8 (1): 980. 10.1038/s42003-025-08414-z.

Yamanaka, Naoki, Kim F. Rewitz, and Michael B. O’Connor. 2013. ‘Ecdysone Control of Developmental Transitions: Lessons from Drosophila Research’. Annual Review of Entomology 58 (1): 497–516. 10.1146/annurev-ento-120811-153608.

Zhou, Xuguo, Matthew R. Tarver, and Michael E. Scharf. 2006. ‘Hexamerin-Based Regulation of Juvenile Hormone-Dependent Gene Expression Underlies Phenotypic Plasticity in a Social Insect’. Development 134 (3): 601–10. 10.1242/dev.02755.

